## Supplementary figures and images for "Limited and strain-specific transcriptional and growth responses to acquisition of a multidrug resistance plasmid in genetically diverse *Escherichia coli* lineages"

### Figure S1

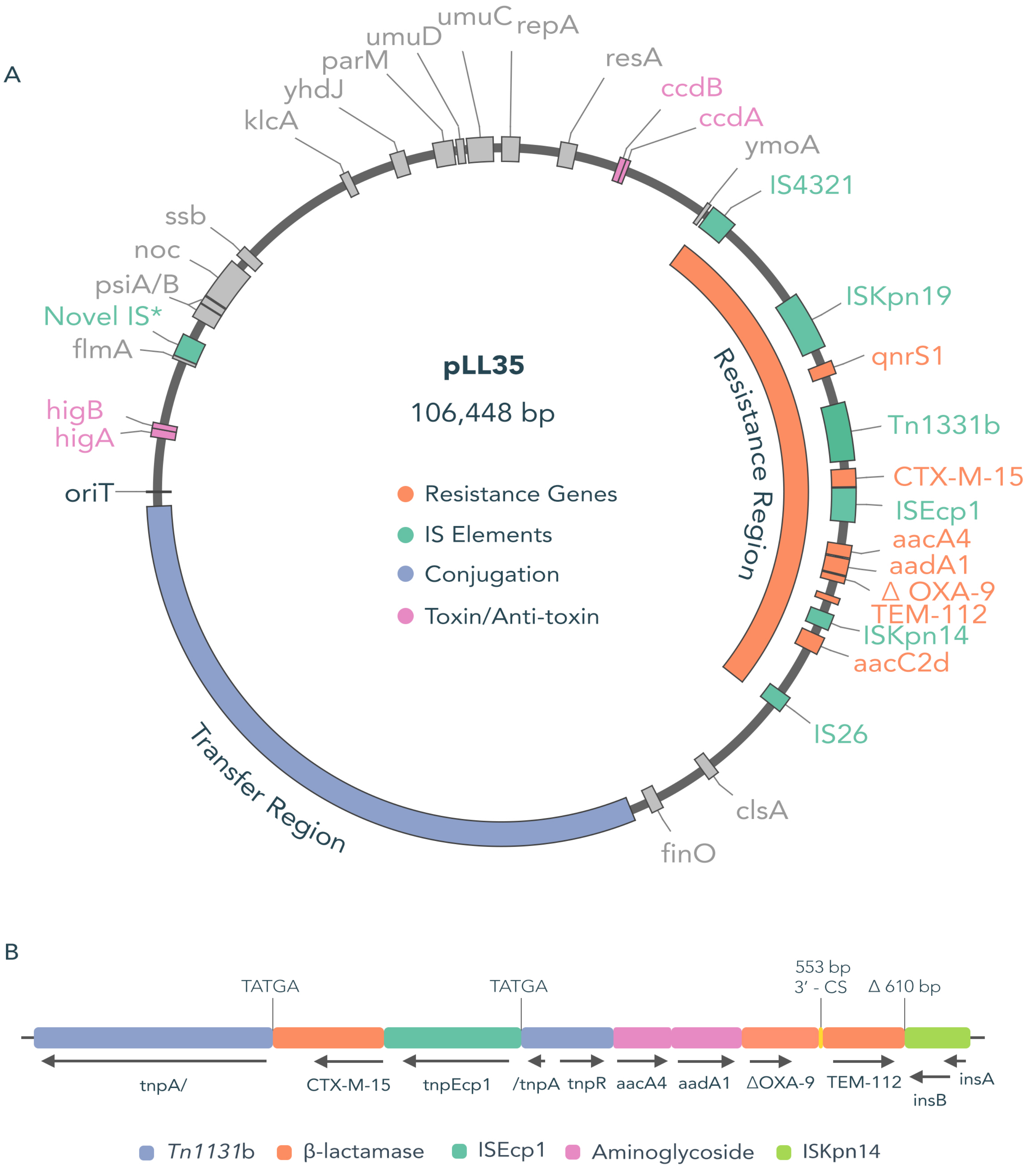

### Figure S2

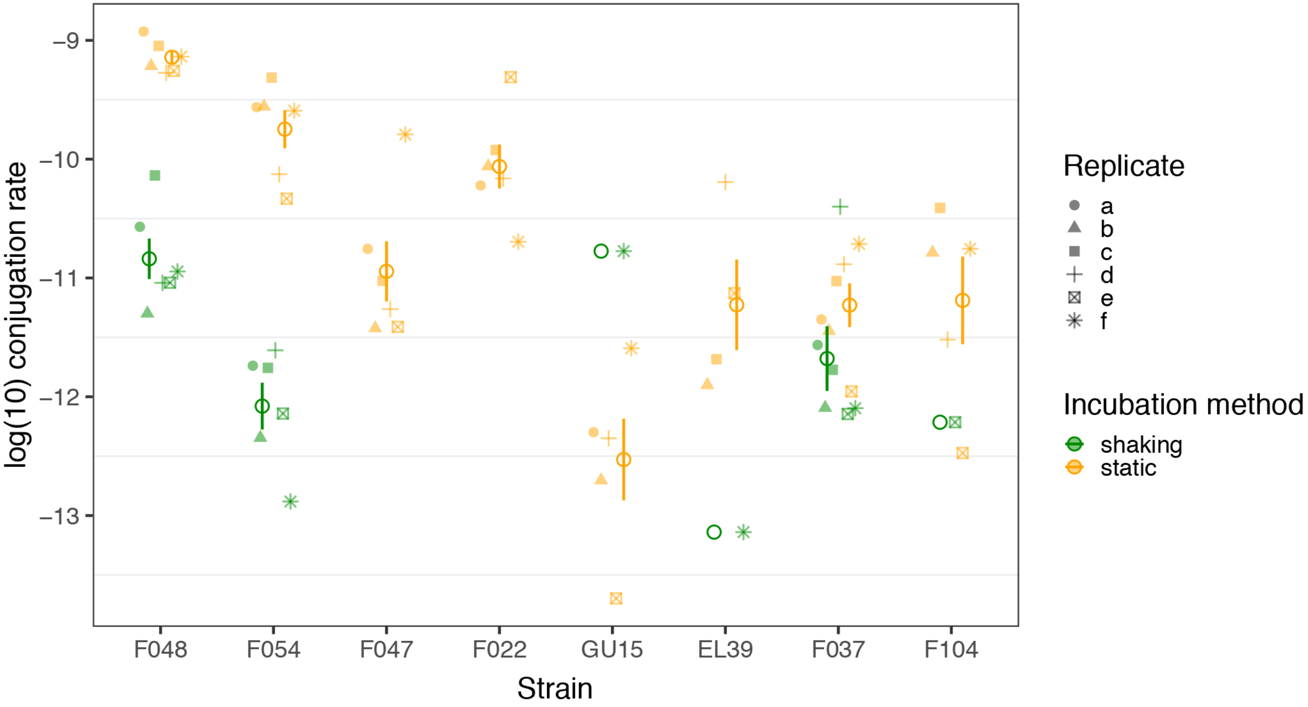

### Figure S3

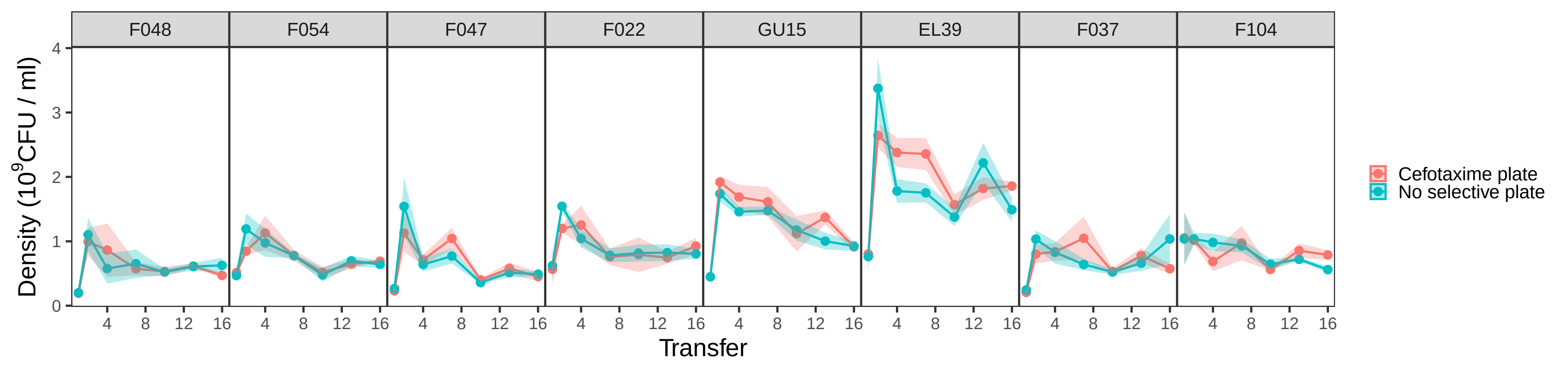

### Figure S4

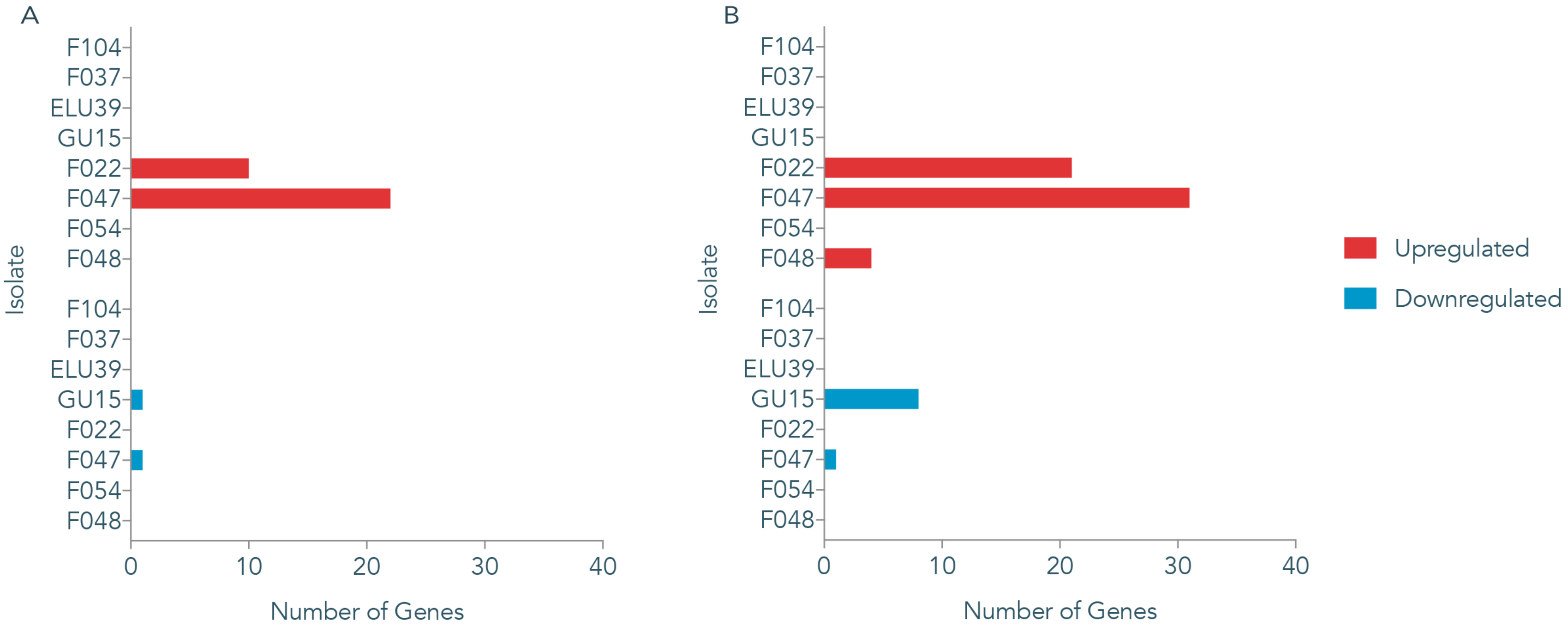

### Figure S6

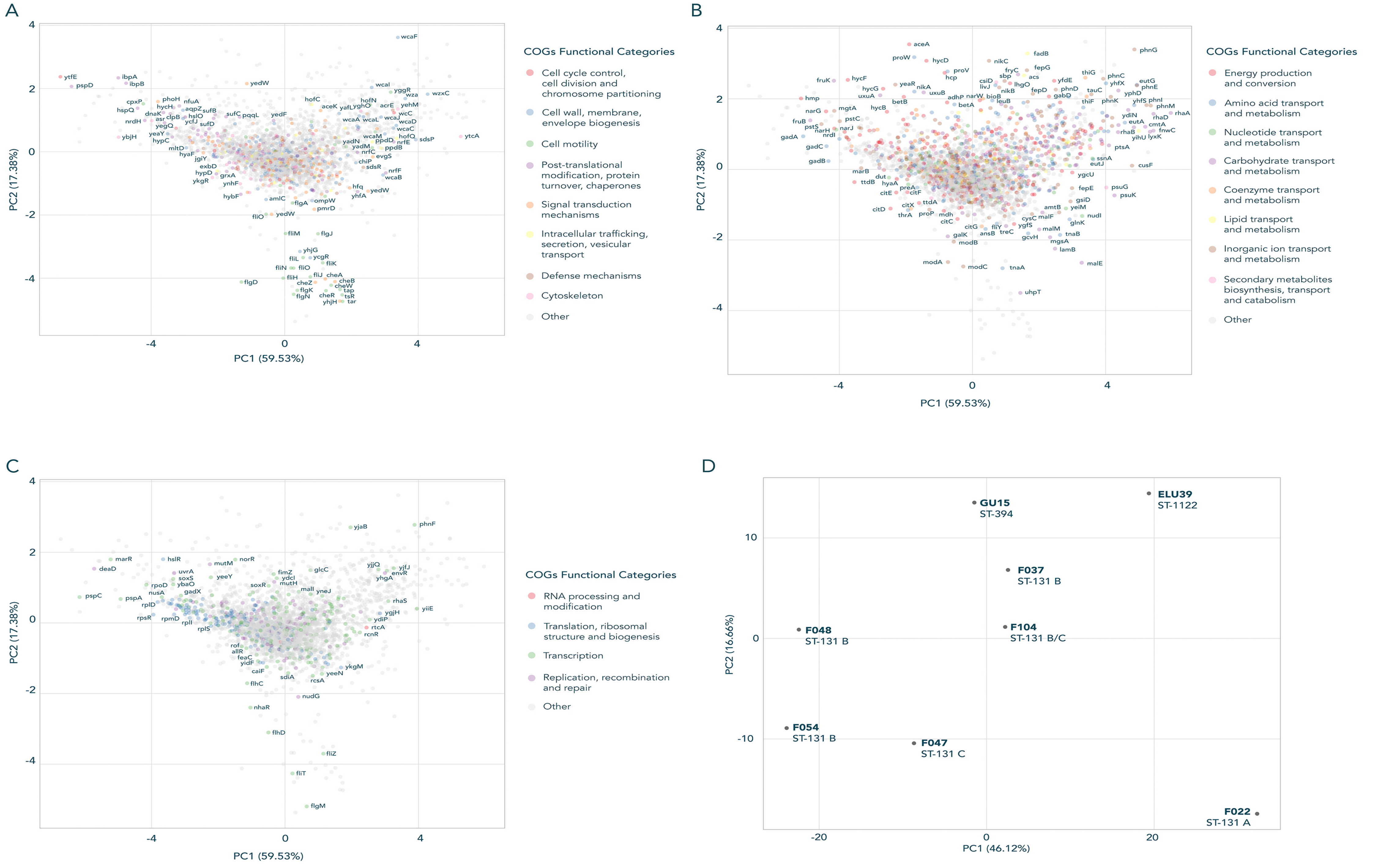

### Fugyre S5

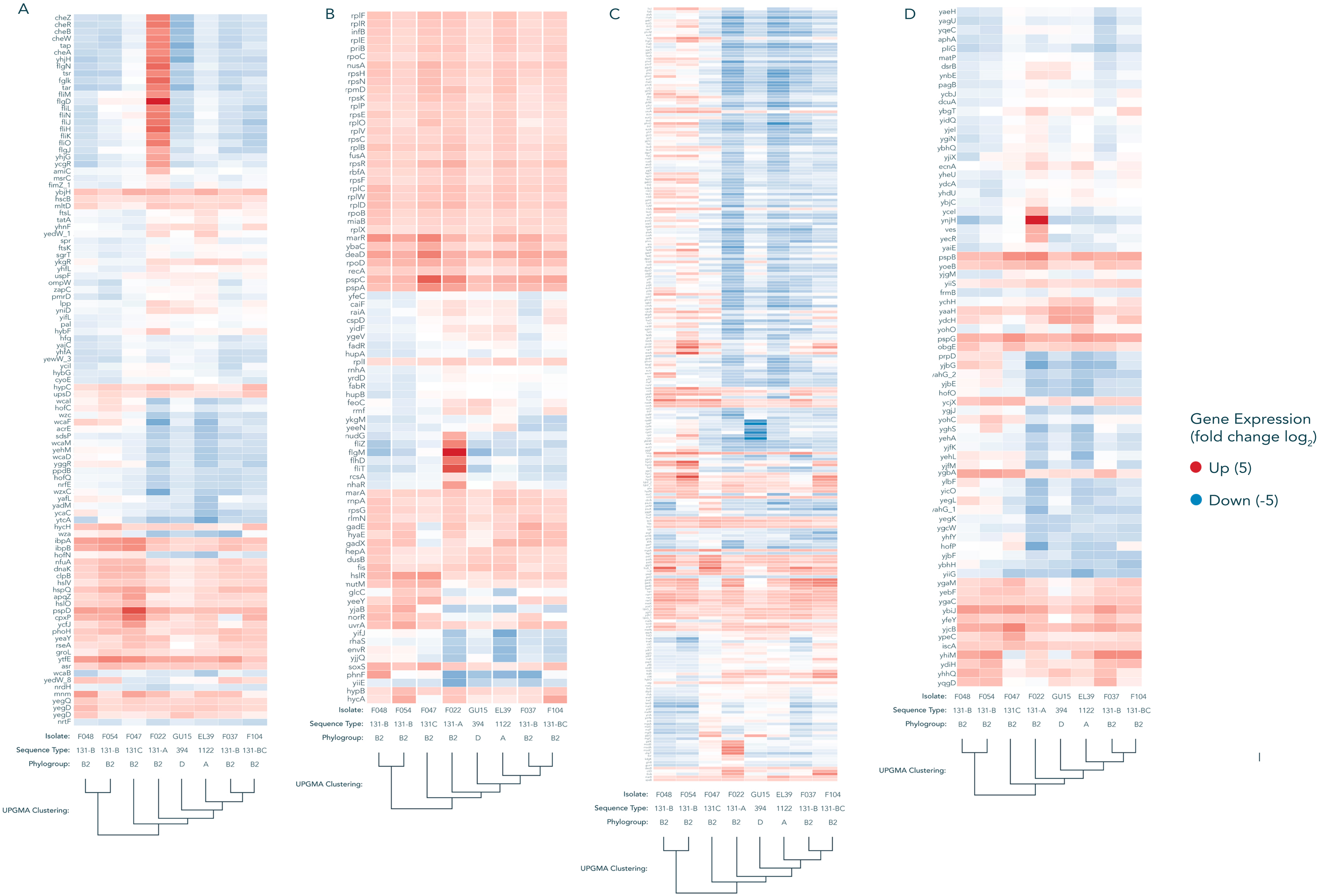
